# Post-embryonic development and aging of the appendicular skeleton in *Ambystoma mexicanum*

**DOI:** 10.1101/2021.03.05.434057

**Authors:** Camilo Riquelme-Guzmán, Maritta Schuez, Alexander Böhm, Dunja Knapp, Sandra Edwards-Jorquera, Alberto S. Ceccarelli, Osvaldo Chara, Martina Rauner, Tatiana Sandoval-Guzmán

**Author notes:** Correspondence Tatiana Sandoval-Guzmán, Technische Universität Dresden, CRTD/Center for Regenerative Therapies TU Dresden, Dresden Germany.

## Abstract

**Background:** The axolotl is a key model to study appendicular regeneration. The limb complexity resembles that of humans in structure and tissue components; however, axolotl limbs develop post-embryonically. In this work, we evaluated the post-embryonic development of the appendicular skeleton and its changes with aging.

**Results:** The juvenile limb skeleton is formed mostly by *Sox9*/*Col1a2* cartilage cells. Ossification of the appendicular skeleton starts when animals reach a length of 10 cm, and cartilage cells are replaced by a primary ossification center, consisting of cortical bone and an adipocyte-filled marrow cavity. Vascularization is associated with the ossification center and the marrow cavity formation. We identified the contribution of *Col1a2*-descendants to bone and adipocytes. Moreover, ossification progresses with age towards the epiphyses of long bones. Axolotls are neotenic salamanders, and still ossification remains responsive to L-thyroxine, increasing the rate of bone formation.

**Conclusions:** In axolotls, bone maturation is a continuous process that extends throughout their life. Ossification of the appendicular bones is slow and continues until the complete element is ossified. The cellular components of the appendicular skeleton change accordingly during ossification, creating a heterogenous landscape in each element. The continuous maturation of the bone is accompanied by a continuous body growth.

## Introduction

Urodeles are the only vertebrates with the ability to regenerate a whole limb as adults, therefore they offer the unique opportunity to gain important insights to advance regenerative medicine. The axolotl (*Ambystoma mexicanum*) is a gold standard for studying limb regeneration in vertebrates. The progenitor cells in the blastema, a key structure for epimorphic regeneration, are a heterogeneous population that remain restricted to their embryonic origin.^1^ Given the cellular heterogeneity of the blastema, the contribution of specific cell populations during regeneration has been the focus of significant research in the last decade. An interesting exception is the skeleton: cells embedded in the skeletal matrix do not participate in regeneration; instead, the skeleton is restored by periskeletal cells and dermal fibroblasts.^2–4^

While mammals are born with partially ossified bones that will conclude postnatally, ossification in most amphibians is coupled with metamorphosis.^5–7^ However, axolotls remain neotenic and, hence, they maintain juvenile features throughout life and only rarely undergo metamorphosis. Their adulthood is marked by reaching sexual maturity, a slowing growth rate, and by ossification of the appendicular skeleton.^8, 9^ The physiological context in which appendicular ossification occurs in axolotls as well as the cellular transitions within the skeleton are unclear. In the anuran amphibian, *Xenopus laevis*, the extent of ossification negatively correlates with the regenerative potential of the limb. During metamorphosis and as bone ossifies, the regenerative capacity declines in the ossifying areas, while amputation at the cartilaginous joints still regenerates.^10^ Limb regeneration in salamanders remains a feature of adult animals; however, regeneration also declines with age and after metamorphosis.^11^ It remains unclear if the correlation between appendicular ossification and decreased regeneration holds true for the axolotl, and more importantly, if the regeneration mechanisms studied so far are replicated when the cellular and extracellular matrix landscapes change in the skeleton as they age. In this study, we analyzed larvae, adult and aged axolotls to document the transition of the appendicular skeleton from a cartilaginous to ossified skeleton.

In general, appendicular bones develop by endochondral ossification, a process where a cartilage anlage is replaced by bone.^12, 13^ During development, mesenchymal progenitors condense and form a cartilage primordium which expands by proliferation of chondrocytes. Cells located in the central diaphysis differentiate into hypertrophic chondrocytes (HCs), which subsequently induce the recruitment of blood vessels, osteoclasts and osteoblasts, giving rise to the primary ossification center. Within the diaphysis, the cartilage matrix is degraded, osteoblasts replace the tissue with cortical bone and a marrow cavity is formed. Simultaneously, osteoblasts located in the perichondrium form a bone collar around the diaphysis and the periosteum is established. Many HCs undergo apoptosis during endochondral ossification; however, several studies have shown a partial contribution from HCs to bone formation through transdifferentiation into osteoblasts.^14–16^ Moreover, Giovannone *et al.* showed that HCs can also transdifferentiate into adipocytes in the ceratohyal bone in zebrafish, highlighting the importance and participation of HCs in the ossification process. In axolotls, however, endochondral ossification is a post-embryonic process. Whether it is driven by the hormonal changes of sexual maturity, environmental cues or body mass, is not yet determined.

In this work, we evaluated the post-embryonic development of the appendicular skeleton of axolotls, from larvae to aged adults. We analyzed the morphological and cellular changes in the zeugopodial elements, the radius and ulna, and we identified the timing in which ossification of those elements occurred. Moreover, we examined the cellular landscape of the radius and the possible contribution of chondrocytes to bone formation in ulnas. Finally, we observed in adult and aged animals (5-to 20-years) a continuous ossification in all limb skeletal elements. Hence, our work provides a detailed view of how the appendicular skeleton, particularly the zeugopod, matures from a cartilage anlage towards a bone.

## Results

### Axolotls transition from a rapid to a slower growth phase at about 10 months of age

Few studies include aged animals beyond the well-determined larval stages^17^ and the neoteny of the axolotl raises controversy on the limit of its growth. In this study, we used animals grown in fully standardized conditions^18^ and we defined their appendicular development in relation to their maturation stage.

Using the snout to tail (ST) and snout to vent (SV) lengths, we recorded the size and age in 220 axolotls. ST and SV lengths are highly correlated (Pearson coefficient *r* = 0.9937, *p* = 3.1 10^-209^, Fig. 1A) and their normograms presented a similar trend: a rapid growth phase followed by a slower growth phase (Fig. 1B, 1C, dots). To test whether there is a transition between two subsequent growth modes, we followed a previously reported approach^19, 20^ to determine the border separating two spatial regions within the anterior-posterior axis of the axolotl spinal cord during regeneration (see methods section). We fitted the ST and SV normograms with a two-line mathematical model, assuming two subsequent linear growths separated by a transition age (Fig. 1B, 1C, continuous line). We estimated the two-line transition age in (*a_t_^TL^*): 19.87 ± 0.07 and 19.00 ± 0.07 months for ST and SV normograms, respectively (where the errors were estimated by bootstrapping). The slope of the first line (*m_1_*) was higher than the slope of the second line (*m_2_*) for the ST normogram (*m_1_* = 1.144 ± 0.003 cm/month *vs*. *m_2_* = 0.0325 ± 0.0006 cm/month, no overlapping within three times the errors) as well as for the SV normograms (*m_1_* = 0.618 ± 0.002 cm/month *vs*. *m_2_* = 0.0202 ± 0.0004 cm/month, no overlapping within three times the errors). Interestingly, the slope of the second line was higher than zero (*i.e*., zero was smaller than the second line slope minus three times its error, both for ST and SV lengths), indicating that after the transition time, the axolotl continues growing.

**Figure 1.**
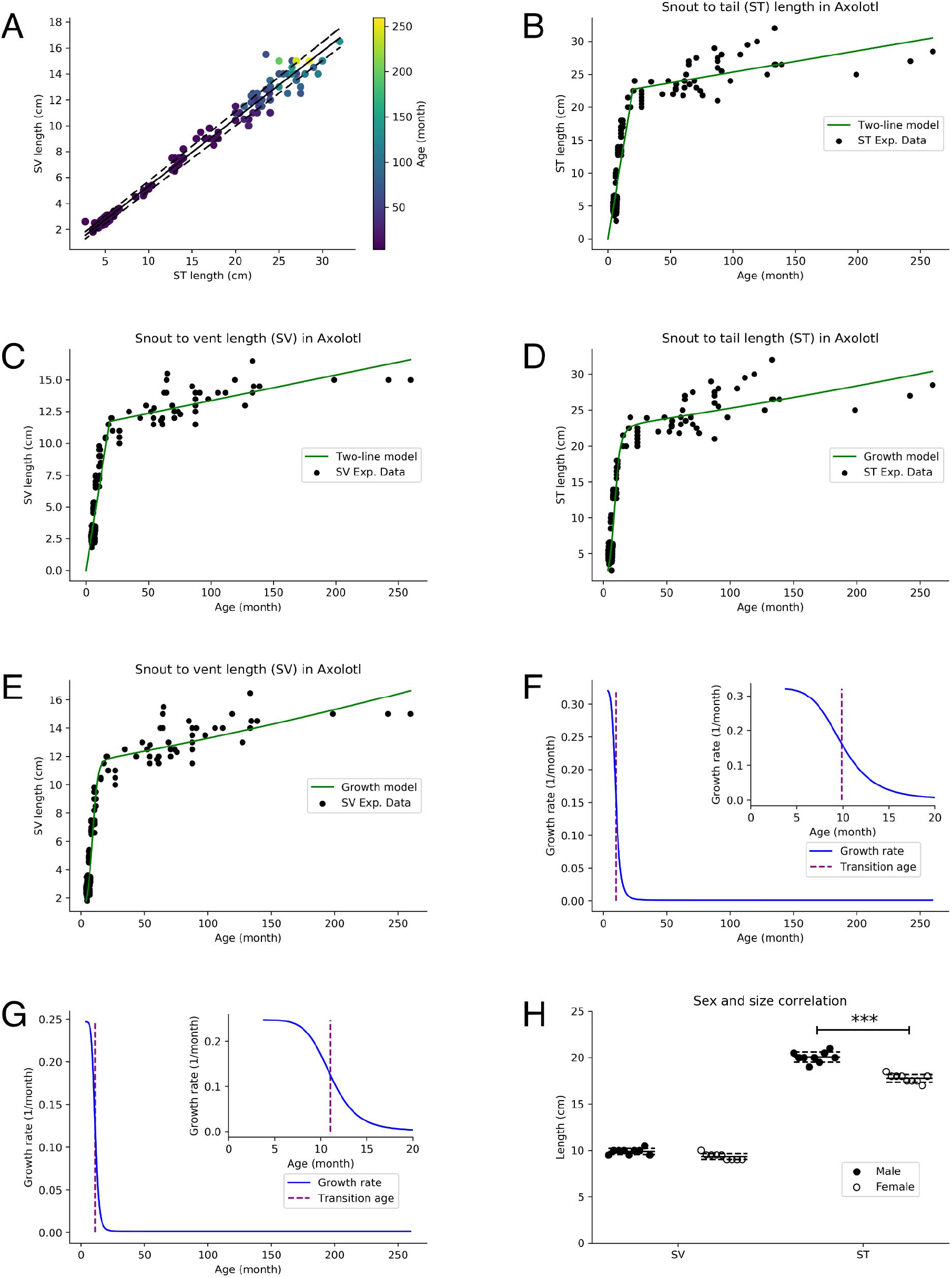
Axolotls transitions from a rapid to a slower growth phase at about 10 months of age. (A) Snout to tail length (ST) linearly correlates snout to vent length (SV) in n = 220 axolotls of different ages (dots). Color code represents age in months. The line shows a linear regression to the experimental data. (B - C) Normogram of ST (B) and SV (C) lengths versus axolotl age (dots, same data shown in A). A two-line model was fitted to the experimental data (the continuous lines depict the best-fitting simulation). (D - E) A growth model was fitted to the experimental ST (D) and SV (E) lengths (dots, same data shown in A, B and C; the continuous lines depict the best-fitting simulation). (F - G) Time course of the Growth rate predicted by the growth model fitted to the ST (F) and SV (G). Insets show magnification of the age near the transition age (vertical lines) of 9.87 ± 0.05 and 11.07 ± 0.05 months for the ST (F) and SV (G) lengths respectively. (H) Quantification of ST and SV lengths in cm in 1-year-old siblings as a function of its sex. (*n* = 18, 9 males and 9 females. *** *p* < 0.001, Student’s *t* test).

To mechanistically address the descriptive results obtained with the two-line model, we tested whether the normograms could be recapitulated by a linear growth model assuming a time-dependent growth rate (see methods section). In particular, we assumed that the growth rate follows an age-dependent Hill-expression. The model successfully fit both normograms (Fig. 1D, 1E) and allowed us to predict the age-dependent growth rate, a proxy for the global proliferation rate in the axolotl (Fig. 1F, 1G). The model allows us to predict the growth rate for animals of old ages, which resulted in (1/month): 0.00116 ± 0.00003 and 0.00142 ± 0.00002 for ST and SV lengths, respectively, both being higher than zero, in agreement with the result obtained with the two-line model. More importantly, we determined two subsequent growth rate regimes separated by the growth rate transition age (*a_t_^GR^*)given by the inflexion point, estimated as 9.87 ± 0.05 and 11.07 ± 0.05 months for the ST and SV lengths, respectively. Hence, our analysis suggests that the two-line transition age of approximately 20 months is an emergent phenomenon resulting from the growth rate transition age of about 10 months.

The biological context of this transition relates to animals reaching sexual maturity. We documented animal growth in the context of sexual maturity in our axolotl colony. Remarkably, we found that the first signs of secondary sexual features in males appear around 10 months of age, and by 12 months these features are fully developed. However, a successful mating starts only between 11 and 12 months of age. Females develop the first secondary sexual features slightly later than males (between 10-11 months of age) but like males, by 12 months these features are fully developed. Fertility is first successful in females between 12-14 months of age. Hence, the agreement between the transition time separating the faster from the following slower growth rate and the age of appearance of the secondary sexual features suggests a probable causal link.

Of note, in 1-year-old siblings, ST was significantly longer in males than females (males: 20.06 ± 0.6 cm, females: 17.78 ± 0.4 cm, n = 9), while SV remained constant (males: 9.8 ± 0.3 cm, females: 9.3 ± 0.3 cm, n = 9) (Fig. 1H). In 2.5-month-old juvenile larvae siblings, ST did not differ (males: 4.7 ± 0.2 cm, females: 4.7 ± 0.2 cm, n = 12). This indicates that the differences in total length between sexes is due to an increased tail growth in males upon sexual maturation.

### The zeugopodial skeleton is progressively ossified with growth

We evaluated the post-embryonic development of the appendicular skeleton, particularly the long bones from the zeugopod. We used animals ranging from ST 4 to 20 cm, which represent axolotls from a juvenile stage to adulthood. Using alcian blue/alizarin red staining, we first evaluated the broad morphological changes in the skeleton. Alizarin red is a calcium-binding dye that, combined with alcian blue (which stains polysaccharides in cartilage), is routinely used to distinguish bone/cartilage. We detected alizarin red staining in appendicular skeletal elements, starting in axolotls with a ST around 10-12 cm, and expanding along the diaphysis to both ends as the animals grew (Fig. 2A). Alizarin red is detected in a proximodistal developmental progression, from stylopod, to zeugopod and finally autopod. The epiphyses of these elements remained cartilaginous in ST 20 cm animals. It is important to note that alizarin red also labels mineralized cartilage, which can be seen *in vivo* in axolotls by fluorescence imaging (Fig. 2B), hence it does not accurately distinguish the transition from calcification to ossification. To accurately detect and quantify ossified bone volume and trace the start of the ossification process, we analyzed different limbs by µCT scan. Ossification was not detectable in ST 10 cm animals. The first appearance of an ossified ring around both the radius and ulna was observed in animals with a ST of 12 cm (Fig. 2C). In animals bigger than 12 cm, ossification extended longitudinally to both ends of the bone. Interestingly, the ossification did not occur homogenously, but rather with a porous appearance, resembling the woven-like bone seen in mammalian bone development. In ST 25 cm animals, this porosity disappeared and the tissue became uniformly ossified (Fig. 2D), forming compact bone. Ossification extended to the epiphyses of the bone, as observed in the widening ends. Quantification of the bone volumes (Fig. 2E) are not significantly different between bones from ST 12 and 14 cm animals, or between ST 14 and 16 cm ones. However, between ST 16 cm and 20 cm, a 4-fold increase in bone volume was observed for the radius, and 7-fold for the ulna. Moreover, radius volume was significantly higher than ulna (1.4 times increase) in ST 20 cm axolotls. Even though ossification started when animals had a ST of 12 cm, there is variability between siblings, especially at a ST of 14 cm (Fig. 2E). Whether this variability results from sexual dimorphism or simply interindividual variation is yet to be determined.

**Figure 2:**
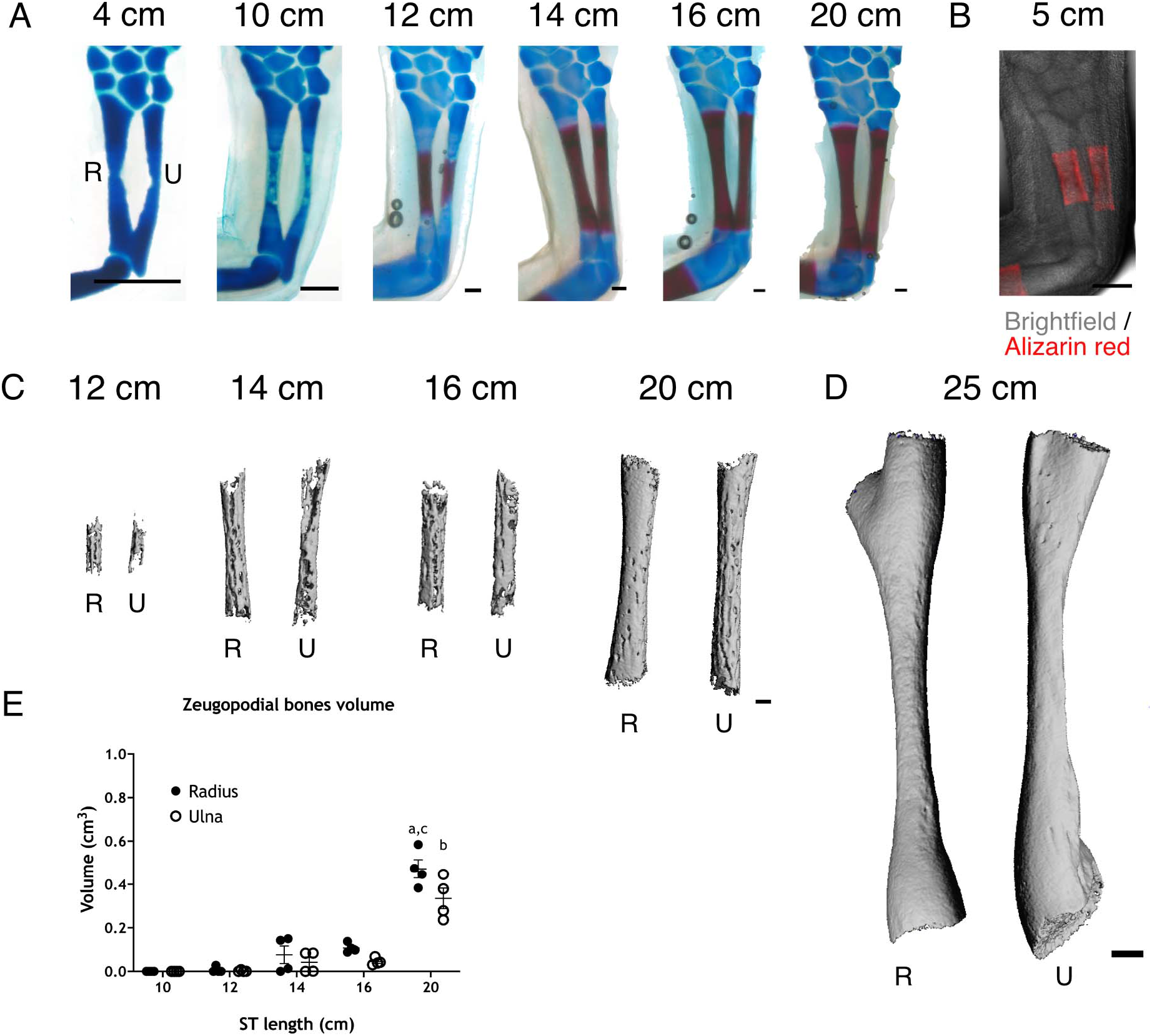
The zeugopodial skeleton is progressively ossified with growth. (A) Alcian blue/alizarin red staining of limbs from different ST animals (n = 4). R: radius, U: ulna. Scale bar: 500 µm. (B) *In vivo* alizarin red staining of axolotl ST 5 cm. Scale bar: 500 µm (C) 3D reconstructions from µCT scan for zeugopodial bones from different ST animals (n = 4). All bones are scaled. Scale bar: 250 µm. (D) 3D reconstruction from µCT scan for zeugopodial bones from ST 25 cm axolotl (n = 1). Bones scaled to (C). Scale bar: 500 µm. (E) Quantification of zeugopodial bones volume (cm^3^) from different ST animals. (n = 4 per size. a: radius ST 20 cm compared to radius from ST 10, 12, 14 and 16 cm. b: ulna ST 20 cm versus ulna from ST 10, 12, 14 and 16 cm. c: radius ST 20 cm versus ulna ST 20 cm. a, b: p < 0.001, c: p < 0.05, Tukey’s multiple comparisons test).

### Zeugopodial bones are vascularized and the marrow cavity is filled with adipocytes

Changes of the skeletal cellular landscape accompany the transition from a cartilage anlage to bone. We assessed these changes in isolated radii from ST 6 and 12 cm axolotls, which corresponded to radii before and during ossification. We performed H&E staining on paraffin sections and confirmed that the radius was exclusively composed of chondrocytes in ST 6 cm animals (Fig. 3A, left panel). We identified a resting zone (RZ), filled with rounded chondrocytes, a proliferative zone (PZ), characterized by flattened chondrocytes, and a hypertrophic zone (HZ), where chondrocyte differentiation leads to an increase in cellular volume. Interestingly, these zones were not uniformly organized, as opposed to the isotropic cellular distribution of mammalian growth plates. The lack of an organized cellular distribution has been previously observed in another salamander model.^7^ Throughout the juvenile stages, a calcification ring surrounds the perichondral region of the HZ (Fig. 3A, arrowhead Fig. 3B). We have observed that the first calcification ring appears early in limb development (stage 48) not long after hatching (stage 44), when the humerus is formed and the radius and ulna have just started to form by bifurcation from the humerus distal end (according to staging table^17^).

**Figure 3:**
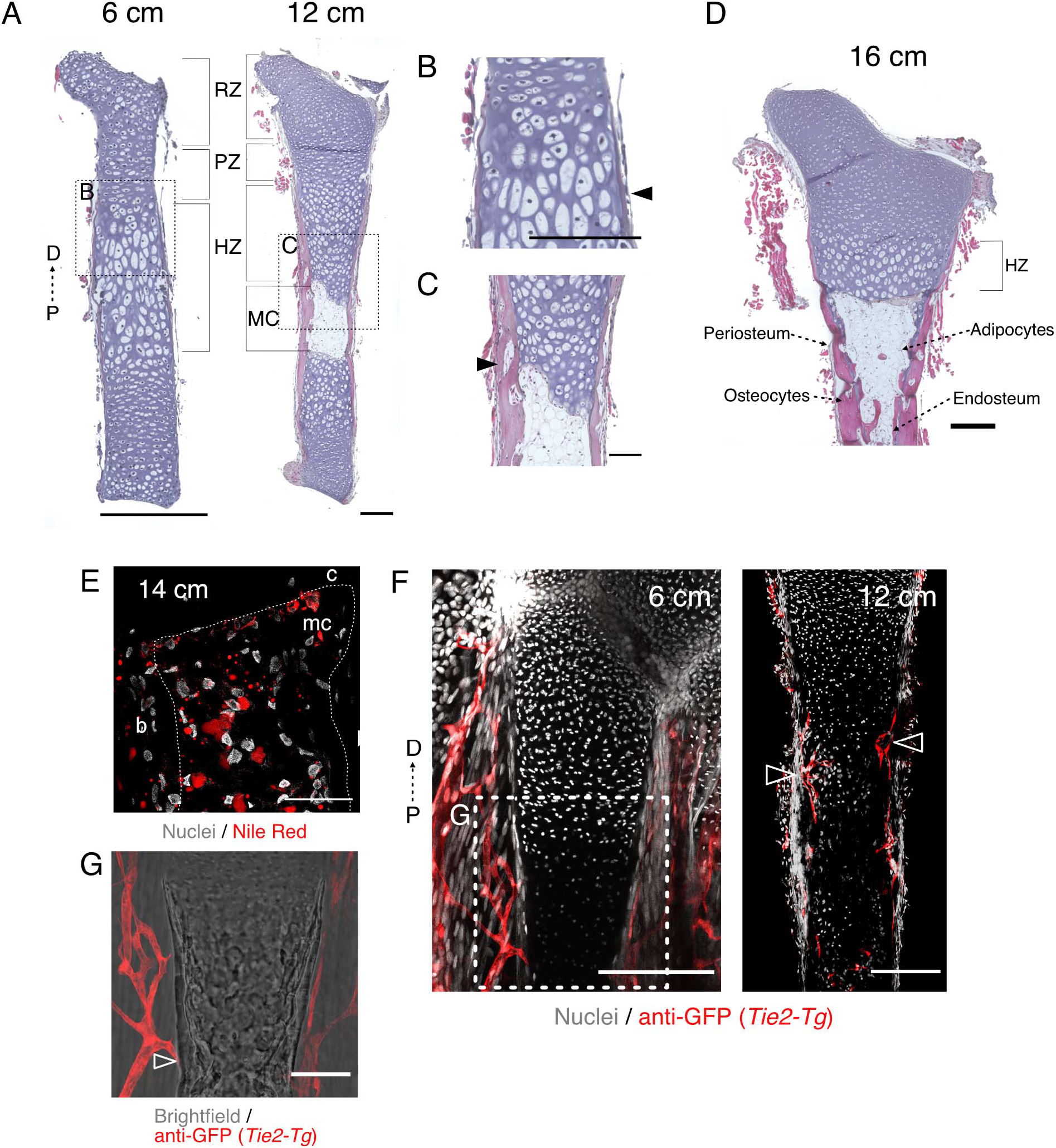
Zeugopodial bones are vascularized and the marrow cavity is filled with adipocytes. (A) H&E staining from radii from axolotls ST 6 and 12 cm (n = 6). RZ: resting zone, PZ: proliferative zone, HZ: hypertrophic zone, MC: marrow cavity, D: distal, P: proximal. Scale bar: 500 µm. (B) Inset from ST 6 cm in (A). Arrowhead: calcified ring around HZ. Scale bar: 250 µm (C) Inset from ST 12 cm in (A). Arrowhead: discontinuous ossification around diaphysis. Scale bar: 250 µm (D) H&E staining in the distal half of the radius from a ST 16 cm axolotl (n = 6). Scale bar: 500 µm. (E) Nile red staining in marrow cavity of a ST 14 cm axolotl (n = 3). c: cartilage, b: bone, mc: marrow cavity. Scale bar: 100 µm. (F) Confocal image of anti-GFP whole mount IF of forelimb or radius from *Tie2-Tg* (red) ST 6 or 12 cm animals, respectively (n = 4). TO-PRO™3 was used for nuclear staining (white). Image represent a maximum intensity projection of 3 images (10 µm interval). Arrowheads: bone vascularization. Scale bar: 250 µm. (G) Inset from *Tie2-Tg* ST 6 cm in (F) with overlapping brightfield image. Arrowhead: blood vessel in contact with calcified cartilage. Scale bar: 100 µm.

Radii from ST 12 cm animals (Fig. 3A, right panel) have a similar cellular distribution of the cartilage region as younger animals. But in contrast to young animals, we observed the formation of a primary ossification center in the diaphysis, which contained a marrow cavity surrounded by cortical bone. The woven-like bone seen on the µCT scan was also observed in sections as discontinuous tissue surrounding the cartilage, with defined edges between gaps (arrowhead Fig. 3C). The bone extended beyond the marrow cavity, surrounding also the HZ. By ST 16 cm, radii stained with H&E showed a further progression in ossification with an expanding marrow cavity. However, cartilage lining subjacent to the bone is still present, suggesting that the transition to bone was not completed. Embedded osteocytes in the cortical bone matrix and cells lining the outer and inner sides of the bone (periosteum and endosteum respectively) were observed (Fig. 3D). In both ST 12 and 16 cm, the marrow cavity was filled with adipocytes, which were both identified by their morphology and by Nile red staining (Fig. 3E), a lipophilic dye commonly used to identify lipid droplets.

A critical step for bone formation by endochondral ossification is the vascularization of the tissue.^21^ To assess the vascularization of the skeletal elements from ST 12 cm animals, we generated a new transgenic line that expresses EGFP under the control of a *Tie2* promoter and enhancer (*Tie2-Tg*),^22^ labeling blood vessels in the entire animal. We collected limbs from ST 6 cm animals and the radii from ST 12 cm animals and performed whole-mount immunofluorescence against GFP, followed by tissue clearing and confocal imaging. We observed that blood vessels surrounded the skeletal element of ST 6 cm animals, but the cartilage was not vascularized (Fig. 3F, left panel). However, we observed some blood vessel protrusions in close contact with the perichondrium in the HZ (arrowhead, Fig. 3G), which could point to the position where the ossification of the skeletal element will start. In the radii of ST 12 cm animals (Fig. 3F, right panel), the mid-diaphysis was found to be vascularized, which correlates with the ossification of the cartilage anlage and formation of the marrow cavity. We did not observe vascularization of the epiphyses, and thus no secondary ossification center formation at this stage.

### *Sox9* marks chondrogenic cells while *Col1a2* marks all skeletal cells

We identified the morphological changes of the zeugopod skeleton during growth; however, the identity of the cells within the skeletal tissue, and their origin, remained unclear. A key transcription factor involved in cartilage development is SOX9^13^. Thus, using a *Sox9-Tg* knock in line, where a T2a self-cleaving peptide followed by mCherry protein is fused to the endogenous SOX9 protein, we evaluated the distribution of mCherry^+^ cells in ulnas from ST 6 and 12 cm animals (Fig. 4A). We confirmed that the expression of mCherry mirrors the real expression of SOX9 by immunofluorescence (Fig. 4E). In tissue sections from ST 6 cm ulnas, most cells were mCherry^+^, while perichondral cells were mCherry^-^ (Fig. 4A, left panel). We observed a similar distribution in ulnas from ST 12 cm animals (Fig. 4A, right panel), and we found no mCherry^+^ cells in the ossified portion of the skeletal element (Fig. 4C). Moreover, using a *Col1a2-Tg* line (which expresses fluorescent protein TFP under the control of the *Col1a2* enhancer, labeling skeletal cells in the axolotl)^4^ we evaluated the distribution of TFP^+^ cells in ulnas from ST 6 and 12 cm animals (Fig. 4B). We observed TFP^+^ cells throughout the skeletal elements in both sizes, labelling both cartilage and bone cells (Fig. 4B, 4D), i.e. chondrocytes, perichondral cells, osteoblasts, osteocytes and periosteal cells.

**Figure 4:**
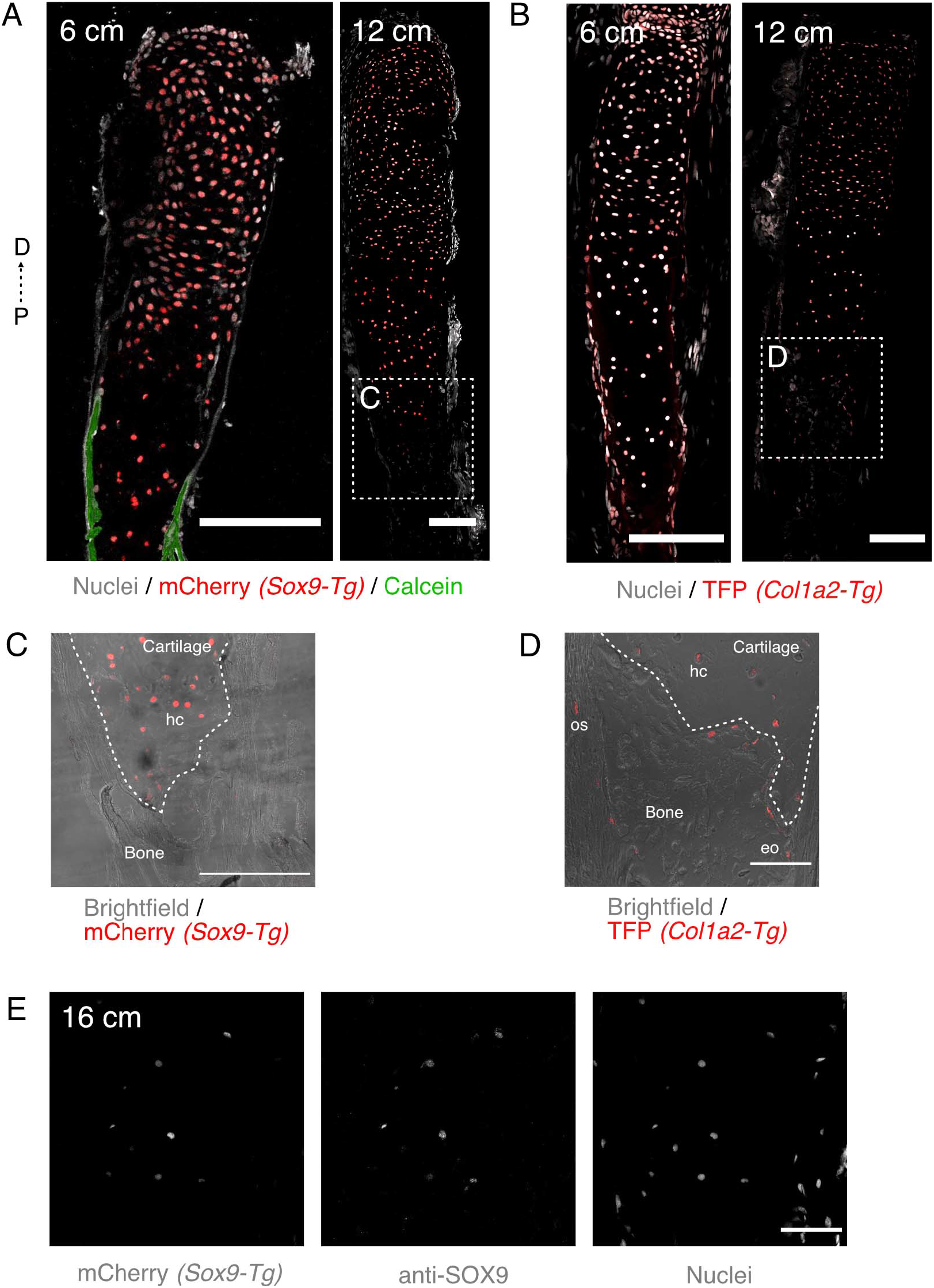
*Sox9* marks chondrogenic cells while *Col1a2* marks all skeletal cells. (A) Confocal image of *Sox9-Tg* (red), ulnas sections from ST 6 and 12 cm axolotls (n = 6). Calcified tissue stained with calcein (green). TO-PRO™3 was used for nuclear staining (white). D: distal, P: proximal. Scale bar: 250 µm. (B) Confocal image of *Col1a2-Tg* (red) ulna sections from ST 6 and 12 cm axolotls (n = 6). TO-PRO™3 was used for nuclear staining (white). Scale bar: 250 µm. (C) Inset from (A) *Sox9-Tg* ST 12 cm. Dotted line shows boundary between cartilage and bone (chondro-osseous junction), hc: hypertrophic chondrocytes. Scale bar: 250 µm. (D) Inset from (B) *Col1a2-Tg* ST 12 cm. os: osteocyte, eo: endosteal cell, hc: hypertrophic chondrocytes. Scale bar: 250 µm. (E) Confocal image of anti-SOX9 IF in ulna sections from *Sox9-Tg* in ST 16 cm axolotl (n = 3). SYTOX™ Green was used for nuclear staining. Scale bar: 250 µm.

### OCN is expressed in bone cells and in some hypertrophic chondrocytes

An unambiguous confirmation of ossification is the identification of mature osteoblasts in the bone matrix and bone lining cells. To that end, we used immunodetection of osteocalcin (anti-OCN). As expected, no labeling was observed in ulnas from ST 6 cm (Fig. 5A, left panel), which further demonstrated the absence of bone tissue in juvenile larvae. In ST 12 cm animals, we observed a broad OCN labeling of the bone ECM (Fig. 5A, right panel). We identified OCN^+^ cells embedded in the bone matrix and in the periosteum (arrowheads, Fig. 5B). Interestingly, OCN^+^ cells were also identified in the lower HZ of the chondro-osseous junction of the mid-diaphysis. These cells were mCherry^+^, identified by our *Sox9-Tg* line (solid arrowheads Fig. 5B, 5C), and SOX9^+^, identified by sequential double immunofluorescence (arrowheads Fig. 5D). Previous studies have correlated the downregulation of *Sox9* in the HZ with the activation of the osteogenic program, and the persistence of SOX9 with an inhibition of bone-related markers.^23, 24^ The expression of OCN in HCs could point towards an early activation of an osteogenic program in those cells. In that regard, osteocalcin expression has been reported in post-HCs.^25^ Some studies have demonstrated the ability of HCs to transdifferentiate into osteoblasts in mammals and zebrafish (Giovannone et al., 2019; Yang et al., 2014). However, the co-expression of SOX9 and OCN needs to be further explored.

**Figure 5:**
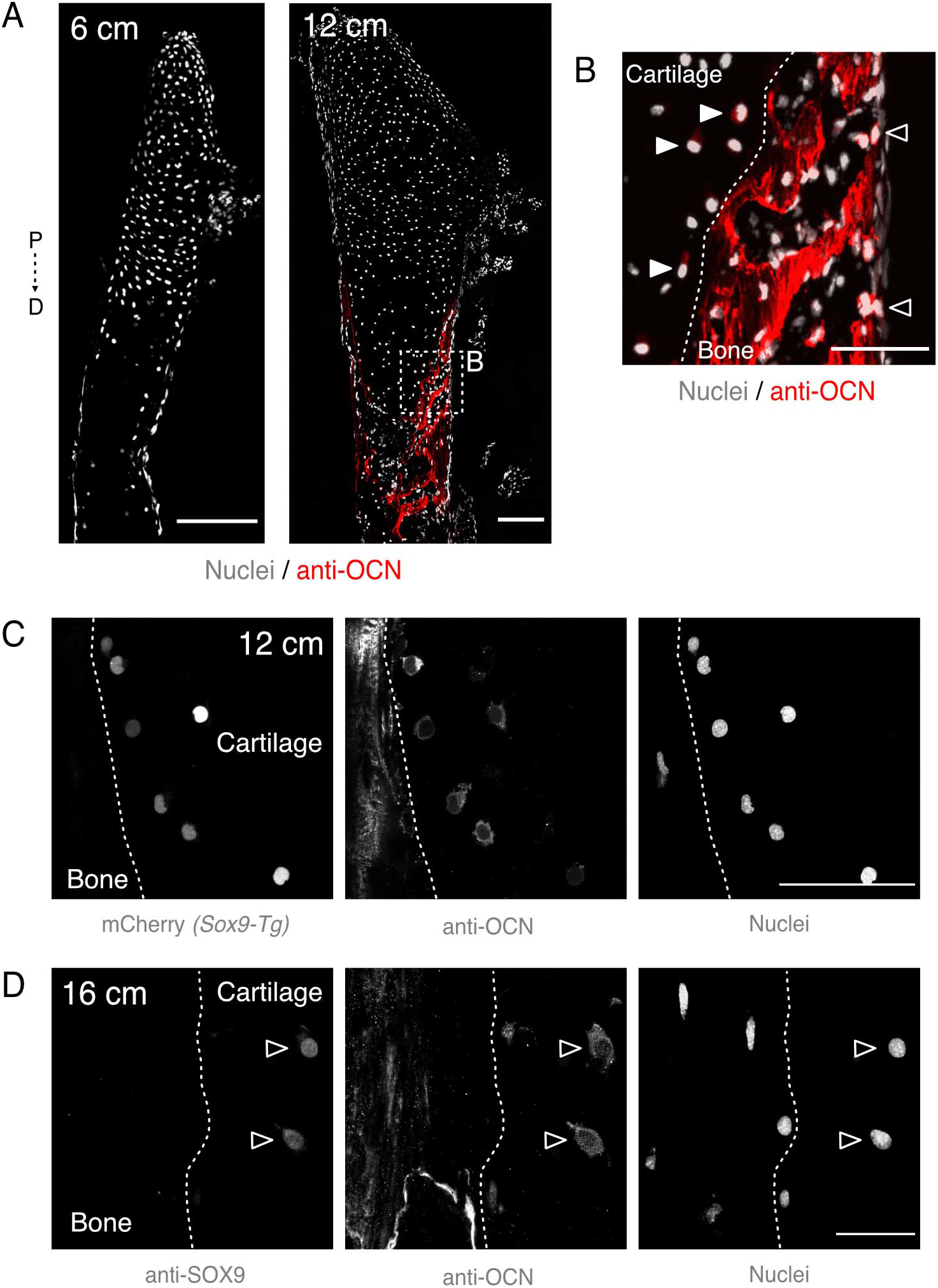
OCN is expressed in bone cells and in some hypertrophic chondrocytes. (A) Apotome image of anti-OCN IF (red) from representative radii sections from ST 6 and 12 cm axolotls (n = 6). SYTOX™ Green was used for nuclear staining (white). D: distal, P: proximal. Scale bar: 250 µm. (B) Inset from (A) anti-OCN IF (red). Dotted line shows boundary between cartilage and bone. Solid arrowhead: hypertrophic chondrocytes. Arrowhead: periosteal cells. Scale bar: 100 µm. (C) Confocal image of anti-OCN IF from radius section from *Sox9-Tg* ST 12 cm (n = 3). SYTOX™ Green was used for nuclear staining. Scale bar: 100 µm. (D) Confocal image of sequential IF for anti-SOX9 and anti-OCN in ST 16 cm axolotls (n = 4). Hoechst 33342 was used for nuclear staining. Image represent a maximum intensity projection of 4 images (2 µm interval). Arrowheads: SOX9^+^/OCN^+^ cells. Scale bar: 50 µm.

### Lineage tracing reveals a contribution of *Col1a2* cells to bone and marrow cells

Our above observations led us to hypothesize that some HCs could transdifferentiate into bone cells in the axolotl. To start exploring this idea, we evaluated the contribution of *Col1a2* chondrocytes to bone formation.

Using the double transgenic *Col1a2xLPCherry-Tg*, we traced the fate of *Col1a2* descendants by inducing the permanent labeling of chondrocytes by tamoxifen-induced conversion of the Cre/Lox reporter cassette in ST 5 cm animals. In these animals the appendicular skeleton is still exclusively cartilaginous. We allowed the animals to grow until ST 14 cm and we collected the ulnas for sectioning and imaging. We observed a great proportion of the cells to be mCherry^+^. They were distributed throughout the cartilage, bone matrix, periosteum, endosteum and in the marrow cavity (Fig. 6A, 6B). Moreover, we assessed whether some of the mCherry^+^ cells were OCN^+^. We found mCherry^+^/OCN^+^ cells in the endosteum and periosteum (arrowheads, Fig. 6C, 6C’, 6C’’), showing that *Col1a2* descendants gave rise, at least partially, to committed osteoblasts in the axolotl zeugopod. However, whether a specific subpopulation of *Col1a2* cells gives rise to all derived cells or if HCs contribute directly to OCN^+^ cells, cannot be answered with this transgenic approach.

**Figure 6:**
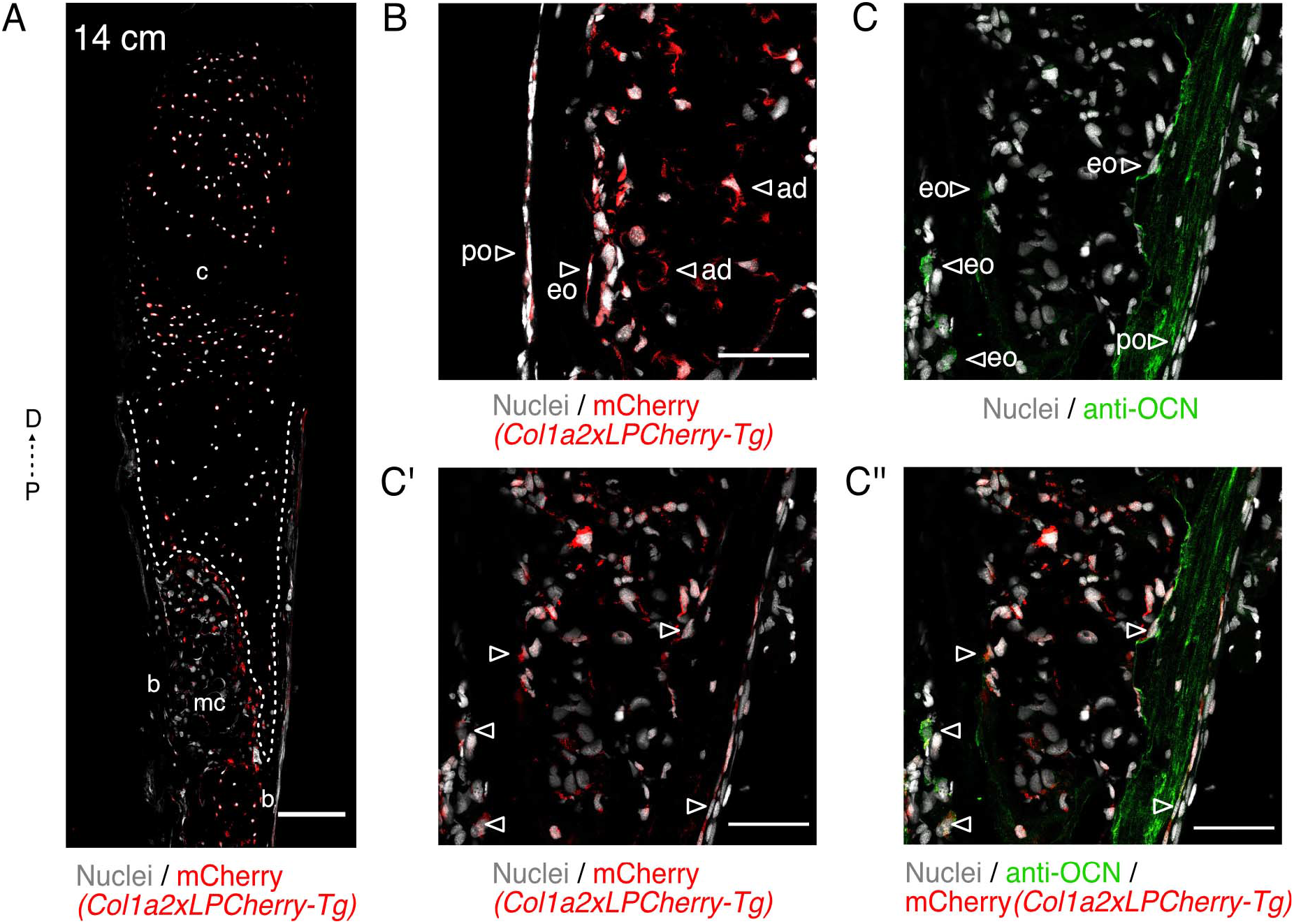
Lineage tracing reveals a contribution of *Col1a2^+^* cells to bone and marrow cells. (A) Confocal image of *Col1a2xLPCherry-Tg* (red) ulna sections from ST 14 cm axolotls (n = 6). SYTOX™ Green was used for nuclear staining (white). Dotted line shows boundary between cartilage and bone, c: cartilage, b: bone, mc: marrow cavity. Scale bar: 250 µm. (B) Confocal image of *Col1a2xLPCherry-Tg* (red) ulna sections from ST 14 cm axolotls (n = 6). SYTOX™ Green was used for nuclear staining (white). Arrowhead: mCherry^+^ cells. po: periosteum, eo: endosteum, mc: marrow cavity. Scale bar: 100 µm. (C-C’’) Confocal image of anti-OCN IF (green) from ulna sections from *Col1a2xLPCherry-Tg* (red) ST 14 cm axolotls (n = 6). SYTOX™ Green was used for nuclear staining (white). Arroheads: mCherry^+^/OCN^+^ cells. Scale bar: 100 µm.

### Bone ossification and marrow cavity expand with age

Axolotls have a life span extending to more than 20-years under controlled laboratory conditions.^8^ However, little is known about bone morphological features in aged animals, although it is commonly assumed that axolotls retain cartilaginous epiphyses and thus there is perennial longitudinal growth of the bones. We collected bones and limbs from multiple 1-, 5-, and 7-year-old axolotls, as well as from two animals that were 20 and 21-years of age. In histological sections from radii (Fig. 7A), we observed an age-related expansion of the marrow cavity towards the epiphyses. Cortical bone was also expanded longitudinally and radially, surrounding most of the skeletal element length, even though some parts of the inner surface of the epiphyses remained cartilaginous in most cases. However, Fig. 7A shows a representative section of the 20-year-old radius, with a partial ossification of the epiphysis (arrowhead), which suggests a continuous ossification of the radius with age. Importantly, we did not identify a secondary ossification center in any of the radii analyzed. Still, we observed the appearance of marrow processes inside the hypertrophic cartilage. These longitudinal projections of the marrow cavity have been described for other salamander models.^26, 27^ In addition, in whole limbs stained with alcian blue/alizarin red, we observed in a 5-year-old animal a calcification in the carpal bone intermedium (Fig. 7B), as well as in 6/8 tarsal bones in our 20-year-old animal (Fig. 7C), and in 4/9 in our 21-year-old specimen. Both aged animals show ossification of the basal commune, central, intermedium and fibular tarsals. This suggests that the short bones in the autopod undergo a late ossification in comparison to the long bones in the zeugopod, but also highlights the continuity of this process in adulthood and aging.

**Figure 7:**
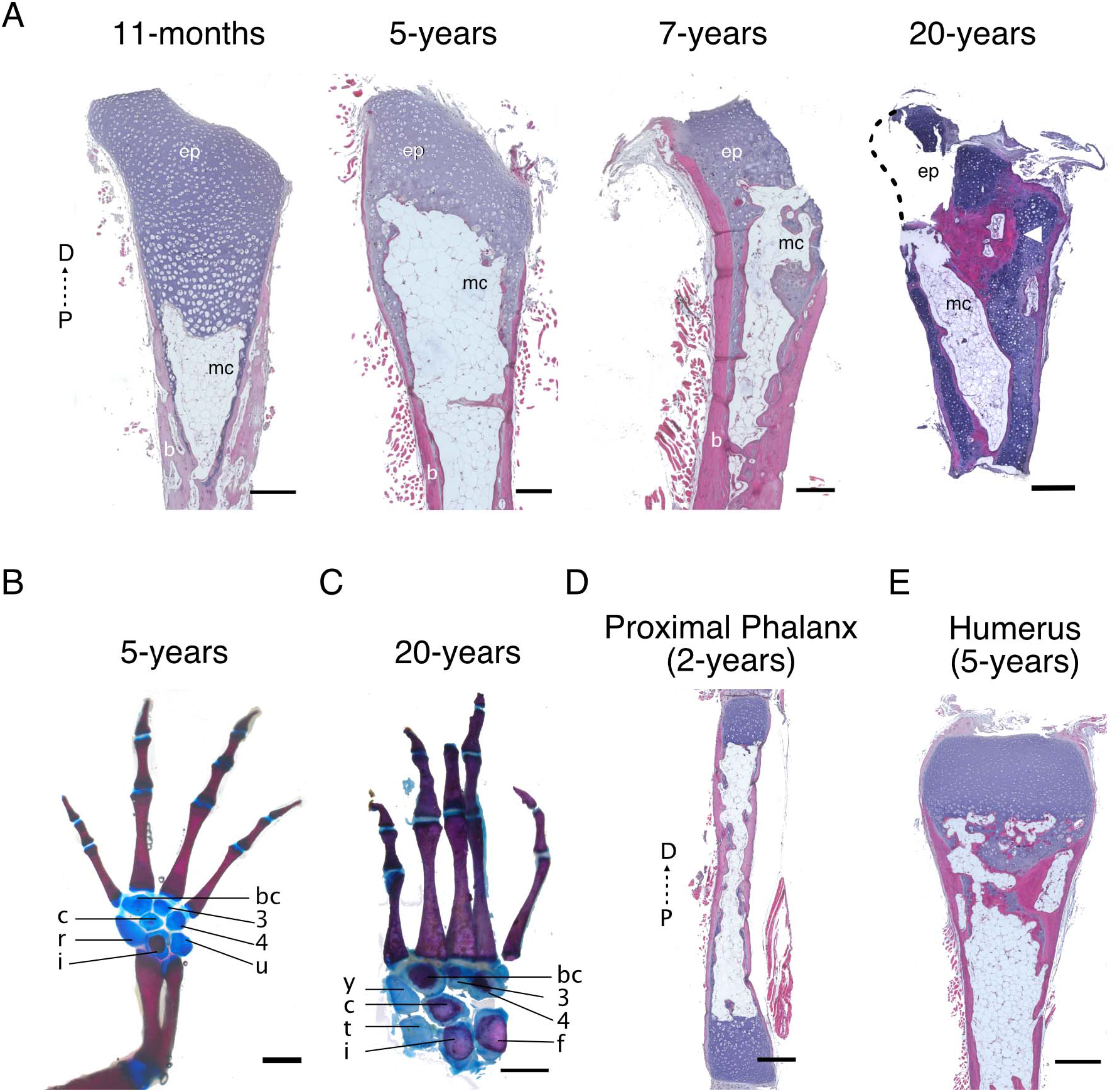
Bone ossification and marrow cavity expand with age. (A) H&E staining of radii sections from different age axolotls (n = 6 for 11-month-old, n = 3 for 5-year-old, n = 5 for 7-year-old, n = 1 for 20-year-old). D: distal, P: proximal, ep: epiphysis, mc: marrow cavity, b: bone. Scale bar: 500 µm. (B) Alcian blue/alizarin red staining of forelimb from a 5-year-old axolotl (n = 3). bc, basale commune; c, centrale; i, intermedium; r, radiale; u, ulnare. Scale bar: 2 mm. (C) Alcian blue/alizarin red staining of foot from a 20-year-old axolotl (n = 1). bc, basale commune; c, centrale; i, intermedium; t, tibiale; f, fibulare; y, element y. Scale bar: 2 mm. (D) H&E staining of proximal phalanx from 2-year-old axolotl (n = 4). Scale bar: 1 mm. (E) H&E staining of humerus from 5-year-old axolotl (n = 2). Scale bar: 1 mm.

In elements from the stylopod and autopod (digits from 2-year-old animals and humeri from 5-year-old animals) (Fig. 7D, 7E respectively), we observed the same features described for long bones in the zeugopod, i.e., an ossified ring of cortical bone around a marrow cavity filled with adipocyte tissue which expanded from the diaphysis towards the epiphyses.

This continuous ossification could be the origin of aberrant phenotypes. In 3 out of 6 7-year-old animals and in the 20-year-old animal, we observed that the zeugopodial bones were fused either in their proximal or distal epiphysis. In one case, the fusion occurred with a heterotopic ossification and cartilage formation around the distal end of the radius.

Our results show a continuous ossification of the skeletal elements even in aged animals. Whether the appendicular skeleton would become completely ossified if an animal lived long enough remains unknown.

### Axolotl metamorphosis accelerates bone ossification

Metamorphosis is a critical transformation in the life cycle of many amphibians. This process allows them to access a vaster territory in their search for food and mating partners. In some anurans and urodeles, ossification of the limb skeleton is associated with the metamorphic transformation.^5, 7^ Although axolotls are neotenic and rarely undergo spontaneous metamorphosis, the administration of exogenous thyroxine can induce it. We evaluated whether the ossification in axolotl is still responsive to the hormonal changes of metamorphosis. If this is the case, metamorphosis should accelerate the ossification process in axolotls. We injected thyroxine in ST 14 cm axolotls, where ossification is already covering the mid-diaphysis. Radii were collected at 35 days post-injection and embedded in paraffin for sectioning and staining. Metamorphic axolotls were slightly smaller than their paedomorph siblings, although this difference was not significant (paedomorph ST: 15.4 ± 0.4 cm, SV: 8.6 ± 0.3, n = 5, metamorphic ST: 14.7 ± 0.3, SV: 8.4 ± 0.5, n = 6). By H&E staining, we did not observe differences in the gross morphology (Fig. 8A) compared to an un-injected animal. These observations are in agreement with a previous study, in which no significant difference was observed in zeugopodial bones when surface and length were compared between a pre-and post-metamorphic axolotl.^28^ However, when compared using µCT scan, we observed an increase in bone volume in metamorphic animals. Both zeugopodial bones had a bigger ossified area, and the ossification pattern was more uniform than the paedomorph bones analyzed (Fig. 8B). Bone volume quantification showed a significant increase of 4 times in radii volume and over 3 times increase for ulnas (Fig. 8C). These results demonstrate that ossification is accelerated in axolotl limbs when metamorphosis is induced, and suggest that the cells respond to intrinsic programs as well as hormonal changes.

**Figure 8:**
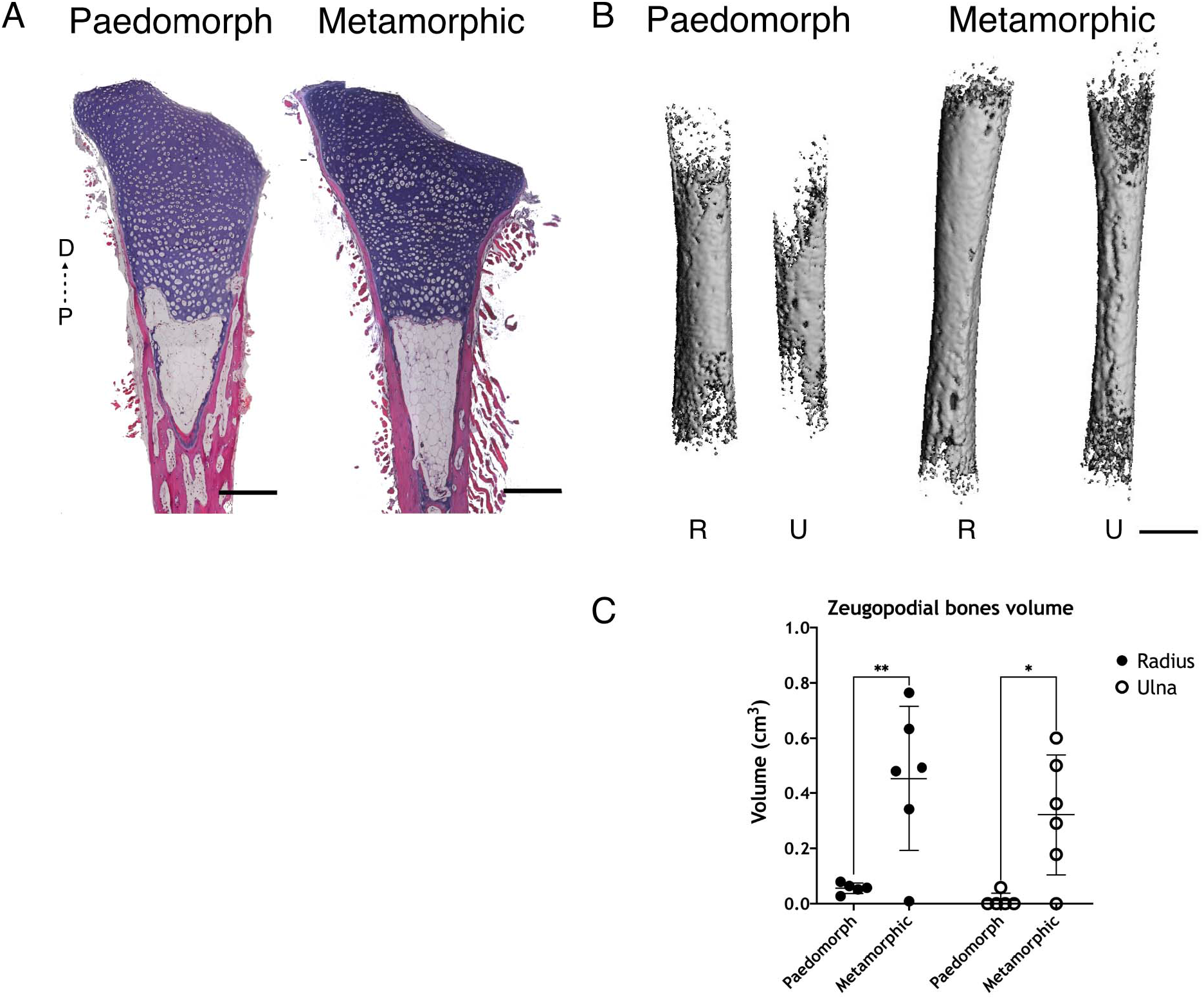
Axolotl metamorphosis accelerates bone ossification. (A) H&E staining from radii sections from paedomorph and metamorphic axolotls (35 dpi) (n = 4). D: distal, P: proximal. Scale bar: 500 µm. (B) 3D reconstructions from µCT scan for zeugopodial bones from paedomorph and metamorphic axolotl (n = 5 for paedomorph zeugopods, n = 6 for metamorphic zeugopods). All bones are scaled. R: radius, U: ulna. Scale bar: 500 µm. (C) Quantification of zeugopodial bones volume (cm^3^) from paedomorph and metamorphic axolotl. (n = 5 for paedomorph zeugopods, n = 6 for metamorphic zeugopods. ** p < 0.01, * p < 0.05, Tukey’s multiple comparisons test).

## Discussion

Appendicular skeletogenesis is considered highly conserved among tetrapods, mostly based on studies in mammals and chicks,^29^ although the appendicular skeleton has undergone species-specific adaptations due to environmental influences. Many species display variations in important cellular and temporal parameters of limb development. Mammals are born with ossified bone, while most amphibians develop their limbs post-embryonically. The transition of a cartilaginous appendicular skeleton to a bony skeleton is associated with the hormonal changes during metamorphosis.^30, 31^ Metamorphosis in amphibians is a significant event that precedes sexual maturity, often giving the animals biological advantages, such as in their ability to acquire food or travel to new locations. In neotenic salamanders, like the axolotl, the appendicular cartilage is not only a transitional tissue; larvae live with a calcified cartilage throughout their juvenile period. Here we show that their limb skeleton continuously ossifies (in the absence of metamorphosis) until the external layer of both, epiphysis and diaphysis is completely covered by compact bone. This result contrast the previously proposed idea that the epiphyses remain cartilaginous throughout life,^26, 28^ and that this feature would likely contribute to longitudinal bone growth.^32^

The biological clock triggering skeletal changes remained unexplored, since axolotls rarely change to a terrestrial environment. In this work, we provide a characterization of the maturation of the appendicular skeleton during the life span of the axolotl, considering ossification timing and age-related effects, complementing previous studies in larval stages^33^ and young adults.^32^ In our axolotl colony, animals are grown in fully standardized conditions that allow us to use the total body length to associate it to appendicular changes and biological landmarks such as sexual maturity. During their growth, snout to vent length scales with the snout to tail length, showing a tightly controlled process. Fitting a two-line mathematical model to our normograms of ST and SV lengths evidence an explosive growth of axolotls in approximately the first 20 months of their life, after which growth velocity dramatically diminishes (Fig. 1B, 1C), however never reaching zero. To mechanistically interpret this transition age, we fitted the same normograms to a linear growth model. Our results suggest that the transition age observed in the growth velocities could be an emergent phenomenon caused by a transition in the growth rates taking place at about 10 months of age (Fig. 1D, 1G). Remarkably, around this time, animals become sexually mature and ossification of their appendicular bones starts, but for most of the juvenile phase axolotls present a mineralized ring around the cartilaginous mid-diaphysis. Mineralization of the zeugopodial appendicular skeletal elements was observed as early as developmental stage 48. In contrast, ossification starts around the sexual maturity period, when we can observe the formation of a denser calcified surface and embedded cells with morphological and genetic characteristics of osteoblasts. Moreover, we show that *Col1a2* descendants participate in the ossification process by giving rise to osteogenic cells and marrow cells (likely adipocytes). We report a continuous ossification of zeugopodial bones and marrow cavity expansion with age. However, this ossification is not complete, as we observed a cartilage-bone composite in most of diaphyses analyzed. Short bones, such as tarsals and carpals, undergo a very late ossification when compared to appendicular long bones. Finally, we see an accelerated zeugopodial ossification in metamorphic animals, indicating a responsiveness to exogenous thyroxine.

### Mineralization *versus* ossification

For most of the initial larval stages, the skeleton is cartilaginous with mineral deposition. We have observed calcification of the humeral diaphysis as early as Stage 48, when the radius and ulna had just started to form by bifurcation from the humerus distal end. Nye *et al*. reported mineralization by alizarin red staining at stage 50, when condensations of the metacarpals are already present.^34^ This discrepancy may be accounted for by the detection protocol. While Nye *et al*. used processed samples, we used real-time vital detection. The calcification of appendicular cartilage remains throughout the juvenile phase of the axolotl until the anlage starts to ossify.

In contrast to mineralization, ossification is a tightly regulated process mediated by osteoblast differentiation. In axolotl limbs, we can identify it around the time animals begin to show secondary sexual characteristics. Generally, amphibian metamorphosis precedes sexual maturation, while in neotenic amphibians both sexual maturation and bone ossification occur in the absence of metamorphosis. In salamanders, as in other species, metamorphosis provides adaptive features to change habitat and ensure greater chances of survival. The axolotl and its geographically and phylogenetically related counterpart, *Ambystoma velasci,*^35, 36^ are examples of facultative metamorphs, which means that individuals transform into terrestrial organisms when the environmental conditions of its habitat change. In facultative metamorphs, ossification happens independently of metamorphosis, although here we questioned if metamorphosis would have an effect on ossification. We show that the genetic program to trigger ossification is indeed independent of metamorphosis, but it is still sensitive to exogenous thyroxine to accelerate the ossification process. In *Xenopus laevis* tadpoles lacking thyroid glands, metamorphosis is halted but growth continues and ossification progresses in the hind limb independently of thyroxine.^37^ Together with our data, this suggests that appendicular ossification programs could also be triggered in the absence of L-thyroxine, and that they probably depend on a critical body mass.

In contrast to the fully ossified diaphysis of post-metamorphic limbs in newts (*Pleurodeles waltl*),^7^ the diaphysis of axolotls retains a cartilage anlage subjacent to the cortical bone even in old animals, and also in post-metamorphic zeugopodial and autopodial elements. We think this observation had eluded previous reports because most of them focus on the humerus or humeral-radial joint. We observed in humeri of a 5-, 7- and 20-year-old animals only a modest cartilage layer in the diaphysis. Although in *Xenopus* the extent of ossification correlates with decreased regenerative potential, axolotls can still regenerate their ossified limbs in paedomorph adults and even in induced metamorph animals.^38^ However, both the potential for regeneration and its fidelity are reduced.^11^ Importantly, our results show that ossification in the appendicular skeleton is continuous. Rux *et al.* observed some degree of ossification of the metaphysis in the femur of a 10-year-old axolotl,^39^ which suggested that ossification of the appendicular bones continues throughout life, and here, we have shown that this ossification indeed reaches the epiphyses with time.

### Contribution of *Col1a2* cells during ossification

One crucial difference between axolotl and mammalian bone formation is the differential activation of the *Col1a2* enhancer. In mammals only osteoblasts activate the *Col1a2* enhancer,^40^ while in axolotls, chondrocytes show *Col1a2* enhancer activity and COL1A2 and COL2A1 expression.^4, 41^ We have observed *Col1a2* enhancer activity as the first mesenchymal condensates form in the limb bud (stage 47), and it remains active in cells inside the cartilage of long bones and in HCs at the chondro-osseous junction even in 7-year-old animals (personal observation).

One cell type largely ignored in regeneration is the adipocyte. We suggested a contribution of *Col1a2* descendants to adipocytes in the marrow cavity. Although our *Col1a2-Tg* line also labels the periosteum, its contribution to homeostasis is typically restricted to skeletal cells. Therefore, we speculate that the likely contribution of *Col1a2* descendants to marrow cells, such as adipocytes, is from chondrocytes. This result agrees with work done in zebrafish, in which lineage tracing of chondrocytes has shown the contribution of these cells to the marrow adipocytes,^14^ similar to what we show in this study. Lopez *et al.* have shown that the axolotl bone marrow is nonhematopoietic in young adults (1-year-old),^42^ and our study detects an adipocyte-filled marrow cavity in the long bones of the zeugopod in young and aged animals (Fig. 3A, 3D, 3E, 7A).

### SOX9^+^/OCN^+^ cells

During endochondral ossification, the fate of HCs is considered to be death and subsequent substitution for bone cells. Unexpectedly, we found that HCs at the chondro-osseous junction are SOX9^+^/OCN^+^. OCN is a non-collagenous protein in the bone matrix expressed and secreted by committed osteoblasts. Although murine HCs can become osteoblasts and osteocytes during endochondral bone formation,^14–16^ it is surprising to find cells expressing both SOX9 and OCN. SOX9 is known to block osteoblast differentiation and to maintain columnar chondrocyte proliferation by delaying pre-hypertrophy, but also to direct hypertrophic maturation.^23, 24, 43^ The roles of SOX9 are various, and its expression in committed osteoblasts has not been shown before, which suggests the presence of chondro-osteogenic hybrid cells during the normal transition to the ossified skeleton in axolotl. Chondro-osteogenic hybrids are generally found in reparative instances, such as in jaw regeneration in zebrafish^44^ and following injury in the murine rib,^45^ and may hold an unexplored potential for bone regeneration. In future studies, with transgenic lines labeling specifically HCs, we will be able to conclusively show their contribution to different cell types in the axolotl skeleton.

### Perspectives

The origin of adipocytes during ossification and their main role inside the marrow cavity needs to be further explored in axolotls. The known role of bone marrow adipocytes is to promote hematopoietic stem cell renewal. However, the marrow cavity of axolotls remains devoid of a hematopoietic niche.^42^ We assume that the adipose tissue in the marrow is white fat based on morphology and the fact that brown adipose tissue has not been reported in amphibians.^46^ As such, it is possible that these cells have, instead, an important role in osteogenesis and energy metabolism in the axolotl. Another cell type of interest is the periskeletal cell. McCusker *et al*. has shown that the periskeletal tissue of older ST 25 cm animals is multilayered and more complex than that of ST 6.5 cm larvae.^3^ Importantly, periskeletal tissue is mainly responsible for the regeneration of the skeleton.^2, 3^ The high potential of these cells has not been tested in non-regenerative conditions, and their origin and contribution to the formation of appendicular bone has not been investigated.

### Concluding remarks

Taken together, our data provide evidence that bone remodeling is a continuous process in axolotls, starting from a mineralized cartilaginous skeleton during the larval stage to a slowly transitioning bone that continues to ossify as animals age. This continuous process is accompanied by a continuous animal growth. The bone ossification occurs in the absence of thyroxine or a biomechanical challenge, suggesting instead the existence of a physiological timer triggered either by sexual maturation or body mass. However, this ossification can be accelerated in response to exogenous thyroxine, creating in a short time a bone mass that would normally take longer to form. Our findings add to a growing body of literature indicating that vertebrate skeletal tissues are structurally diverse, and break many of the established mammalian-centric rules of development and maturation. We consider that understanding the development of the appendicular bones is essential to comprehend how limb regeneration is orchestrated in stages where different cell types and different maturation stages are found at the amputation plane.

## Methods

### Animal husbandry, transgenesis, metamorphosis and sample collection

Husbandry and experimental procedures were performed according to the Animal Ethics Committee of the State of Saxony, Germany. Animals used were selected by their size, which is indicated in each experiment individually (snout to tail = ST; snout to vent = SV). Sex determination in juvenile larvae was performed by PCR based on. ^47^

Axolotls (*Ambystoma mexicanum*) husbandry was performed in the CRTD axolotl facility, adapted from Khattak *et al.*^18^ and according to the European Directive 2010/63/EU, Annex III, Table 9.1. Axolotls are kept at 18-19°C in a 12-h light/12-h dark cycle and a room temperature of 20-22°C. Animals up to 2.5 cm SV are housed in individual tanks with a water surface (WS) of 90 cm^2^ and minimum water height (MWH) of 2.5 cm. Axolotls up to 5 cm SV live in tanks with a WS of 180 cm^2^ and MWH of 4.5 cm. Axolotls up to 9 cm SV live in tanks with a WS of 448 cm^2^ and MWH of 8 cm. Axolotl up to 11.5 cm SV live in tanks with a WS of 665 cm^2^ and MWH of 10 cm. Axolotls of 10-15 cm SV are housed in tanks with a WS of 820-1100 cm^2^ with a MWH of 20 cm. Animals up to 7 cm ST are fed daily with live saltwater artemia, starting at 8 cm ST they are fed with small (3mm) axolotl pellets (Aquaterratec, Norgard Ambrock). Adults of 15 cm ST are fed large pellets (4-4.5 mm) twice a week.

White axolotls (*d/d*) were used for most of the experiments. Transgenic lines used included the previously published TgSceI*(Mmus.Col1a2:TFPnls-T2a-ERT2-Cre-ERT2)^emt^* (referred to as *Col1a2-Tg*) and TgSceI*(Mmus.Col1a2:TFPnls-T2a-ERT2-Cre-ERT2;Mmus.CAGGS:lp-GFP-3pA-lp-Cherry)^emt^* (referred to as *Col1a2xLPCherry-Tg*).^4^ We generated the transgenic lines C-Ti^t/+^*(Sox9:Sox9-T2a-mCherry)^emt^* (referred as *Sox9-Tg*) and TgTol2*(Mmus.Tie2:EGFP)*^tsg^ (referred to as *Tie2-Tg*).

For generation of the *Tie2-Tg* transgenic line, we used the plasmid pSPTg.T2FXK (#54), a gift from Thomas Sato (Addgene plasmid # 35963). The *Egfp* coding region was cloned 3° from the promoter together with *Tol2* sequences. Fertilized embryos from *d/d* axolotls were injected with the *Tie2:EGFP* vector and *Tol2* mRNA as previously described.^18^ The *Sox9-Tg* reporter line is a targeted Knock-in line created by CRISPR/Cas9 technology according to the published protocol.^48^ Briefly, a portion of the Sox9 gene including a part of the second (last) intron and the remaining downstream part of the CDS was PCRed from the axolotl genomic DNA and inserted into the vector pGEM-T along with a DNA fragment encoding T2a-Cherry-3xnls. The resulting vector was injected into fertilized eggs along with the CAS9 protein and the gRNA targeting the sequence GGACTGCTGGCGAATGCACC, which is found in the intron sequence within the vector and in the genome. As a result, mCherry fused to the C-terminus of SOX9 and, separated by a T2a self-cleaving peptide, is expressed from the endogenous SOX9 genomic locus.

Metamorphosis was induced in *d/d* animals as previously described.^18^ Briefly, ST 14 cm axolotls were anesthetized prior to intraperitoneal injections with 1.5 µL of L-thyroxine (Sigma, T2376) per gram of bodyweight (stock 1 µg/µL in DMSO). Animals were cleaned regularly and water level was reduced as gills were lost. Only 1 out of 15 animals died during metamorphosis. Tissue was collected at 35 days post injection.

Recombination of *Col1a2xLPCherry-Tg* was induced by keeping animals in separated tanks with water containing (Z)-4-hydroxytamoxifen (4-OHT, Sigma T5648) 2 µM overnight. Animals were then washed and screened two weeks later to ensure conversion.

For tissue collection, animals were anesthetized with 0.01% benzocaine solution. After collection, animals were euthanized by exposing them to a lethal dosage of anesthesia (0.1% benzocaine) for at least 20 min.

### Alcian blue/alizarin red staining

Limbs collected were fixed with formaldehyde 10% at 4°C for 1 week and then washed three times with PBS 1x and dehydrated with serial EtOH washes (25, 50 and 70%). Limbs were either stored at −20°C in EtOH 70% or processed immediately at RT. Epidermis from animals > ST 12 cm was removed. Limbs were stained with alcian blue solution (Sigma A3157; alcian blue 0.0001% in EtOH 60% / glacial acetic acid 40%) for 3 days. Then, samples were rehydrated with EtOH dilution series in H_2_O (80, 70, 50, 25%), before treatment with trypsin 1% in borax 30% for 30 min. Limbs were rinsed with KOH 1% and stained with alizarin red solution (Sigma A5533; alizarin red 0.0001% in KOH 1%) for 3 days. Next, limbs were washed with KOH 1% and cleared in KOH 1% / glycerol 20% overnight. Finally, limbs were dehydrated with EtOH dilution series (25, 50, 70, 90 and 3x 100%) and later transferred into serial washes of glycerol/EtOH (1:3, 1:1 and 3:1). Limbs were stored in 100% glycerol at 4°C.

### µCT scan

Limbs collected were fixed with formaldehyde 10% at 4°C for 1 week and then washed three times with PBS 1x and dehydrated with serial EtOH washes (25, 50 and 70%) and stored in EtOH 70% at −20°C until measurement. Bone volume and microarchitecture were determined by micro-computed tomography (μCT, vivaCT40, ScancoMedical) of zeugopodial bones using pre-defined script 6. The isotropic voxel size was 10.5 μm (70 kVp, 114 μA, 200 ms integration time). Diaphysis scans were done for the whole zeugopod (both elements at the same time) and included 600 – 1000 slices. ScancoMedical protocols were used for all analyses and 3D reconstructions. Threshold used for analyses in Fig. 2 was 220 mg HA/cm^3^, and for Fig. 8 was 150 mg HA/cm^3^.

### Paraffin sectioning and H&E staining

Skeletal elements were isolated and fixed with MEMFa 1x (MOPS 0.1M pH7.4 / EGTA 2mM / MgSO4×7H2O 1mM / 3.7% formaldehyde) overnight at 4°C, washed with PBS 1x and decalcified with EDTA 0.5M pH7.4 at 4°C for 1 to 4 weeks. After decalcification, samples were washed with PBS 1x and dehydrated with serial EtOH washes (25, 50, 70 and x3 100%). Samples were then incubated 3x with Roti®Histol (Carl Roth, 6640) at RT and 4x with paraffin at 65°C in glass containers. After last incubation, samples were embedded in paraffin (Roti®-Plast, Carl Roth, 6642) using plastic containers and stored at RT. Longitudinal sections of 6 µM thickness were obtained.

H&E staining on paraffin sections was performed following the producer recommendations (Sigma, Procedure No. HHS).

### Cryosectioning and immunofluorescence

Skeletal elements were isolated and fixed with MEMFa 1x overnight at 4°C, washed with PBS 1x and decalcified with EDTA at 4°C for 48 hours. Next, the skeletal elements were washed with PBS 1x and incubated overnight with sucrose 30% at 4°C. Samples were embedded in O.C.T. compound (Tissue-Tek, 4583) using plastic molds and frozen with dry ice for 1 hour prior to storage at −20°C. Longitudinal sections of 12 µm thickness were cut and mounted on superfrost slides. Slides were kept at −20°C until processed.

For immunofluorescence, slides were dried at RT for at least 1 hour. Sections were washed 3x with PBS 1x + 0.3% Tx-100 prior to blocking with PBS 1x + 0.3% Tx-100 + 10% normal horse serum for 1 hour. Primary antibody incubation was done in blocking solution overnight at 4°C. Sections were then washed 3x with PBS 1x + 0.3% Tx-100 and incubated with secondary antibody 1:200 and nuclear staining 1:1000 for 2 hours. Finally, sections were washed 3x with PBS 1x + 0.3% Tx-100 and mounted using Mowiol mounting medium (Carl Roth, 0713).

Sequential IF for SOX9/OCN was performed since the host for both antibodies is rabbit. Protocol was as follows: First IF was done using the anti-SOX9 antibody as previously described. Secondary antibody incubation was done using the donkey anti-rabbit, Alexa Fluor 488. After incubation, slides were washed three times with PBS + 0.3% Tx-100 and blocked with PBS + 0.3% Tx-100 + 10% goat serum for 2 hours. Slides were incubated with anti-OCN antibody in blocking solution for 1 hour at RT and then overnight at 4°C. Slides were then washed 3x with PBS + 0.3% Tx-100 and incubated with goat anti-rabbit, Alexa Fluor 647 antibody in blocking solution for 2 hours. Slides were incubated with Hoechst 1:1000 in PBS + 0.3% Tx-100 for 10 min and then washed with PBS three times for 15 min. Slides were mounted using Mowiol mounting medium. The antibodies and nuclear stainings used in this work are listed in table 1.

**Table 1:** Antibodies and nuclear staining.

| Antibody / Nuclear Staining | Dilution | Cat. Number |
| --- | --- | --- |
| Anti-GFP (chicken) | 1:500 | Abcam (ab13970) |
| Anti-OCN (rabbit) | 1:250 | Abcam (ab198228) |
| Anti-SOX9 (rabbit) | 1:200 | Sigma-Aldrich (AB5535) |
| Goat anti-Rabbit, Alexa Fluor 647 | 1:300 | Invitrogen (A-21245) |
| Donkey anti-Rabbit, Alexa Fluor 488 | 1:200 | Invitrogen (A-21206) |
| Goat anti-Chicken, Alexa Fluor 488 | 1:1000 | Invitrogen (A-11039) |
| Hoechst 33342 | 1:1000 | Invitrogen (H3570) |
| TO-PRO™-3 iodide (642/661) | 1:10000 | Invitrogen (T3605) |
| SYTOX™ Green nucleic acid stain | 1:10000 | Invitrogen (S7020) |

### Whole mount immunofluorescence and tissue clearing

Limbs or isolated skeletal elements were fixed with MEMFa 1x overnight at 4°C, washed with PBS 1x and decalcified with EDTA at 4°C for 48 hours. Whole mount immunofluorescence protocol was adapted from.^49^ Briefly, skeletal elements were washed overnight with PBS 1x + 0.3% Tx-100 at RT and then blocked in PBS + 0.3% Tx-100 + 10% DMSO + 5% goat serum for 24 hours at 37°C. Samples were incubated with primary antibody anti-GFP in blocking solution for 3 days at 37°C. Next, samples were briefly rinsed with PBS + 0.3% Tx-100 and then washed 4x for 2 hours at 37°C. Samples were blocked again at 37°C overnight and then incubated with secondary antibody for 2 days at 37°C. After incubation, samples were rinsed with PBS + 0.3% Tx-100 and then washed 2x for 2 hours at 37°C. Nuclear staining was done for 1 hour in washing solution at RT and samples were then washed 4x in PBS 1x for 15 min at RT. Isolated skeletal elements were embedded in agarose 1% before clearing.

For clearing, samples were dehydrated with serial washes of EtOH (25, 50, 70, 100%) for 2 hours each at 4°C. Samples were then incubated overnight in EtOH 100% at 4°C prior to clearing with ethyl cinnamate (Sigma, 112372-100G) at RT for at least 2 hours. Samples were imaged the same day. The antibodies and nuclear stainings used in this work are listed in table 1.

### Microscopy

Alcian blue/alizarin red staining imaging was performed on a Zeiss Discovery.V20 stereomicroscope (Plan S 1.0x). H&E staining imaging was performed on a Zeiss Axio Observer.Z1 inverted microscope (Plan-apochromat 20x/0.8). Immunofluorescence imaging was performed on a Zeiss Axio Observer.Z1 inverted microscope with an ApoTome1 system (Plan-apochromat 10x/0.45) and on a Zeiss confocal laser scanning microscope LSM 780 (Plan-apochromat 10x/0.45 or Plan-apochromat 20x/0.8). Whole mount immunofluorescence imaging was performed on a Zeiss confocal laser scanning microscope LSM780 (Plan apochromat 10x/0.45).

All images were processed using Fiji.^50^ Processing involved selecting regions of interest, merging or splitting channels and improving brightness levels for proper presentation in figures. Maximum intensity projections were done in some confocal images and it is stated in the respective figure’s descriptions. Stitching of tiles was done directly in the acquisition software Zen (Zeiss Microscopy).

### Fitting the two-line model to the experimental normograms

In this study, we fitted the following Two-line model to the normograms of ST and SV lengths:

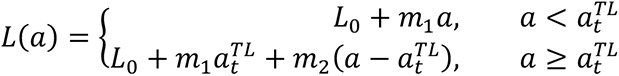

Where, *L*(*a*) is the ST or SV length at age *a*, *L_0_* is the ST or SV length at age zero. The four model free-parameters are *m_1_* is the slope of the first line, *a_t_^TL^* is the transition time of this model and *m_2_* is the slope of the second line.

In our normograms, each value represents a single animal. By fitting the model to the experimental data and using bootstrapping, we estimated the parameter values together with their errors from the ST (table 2) and SV (table 3) lengths:

**Table 2:**
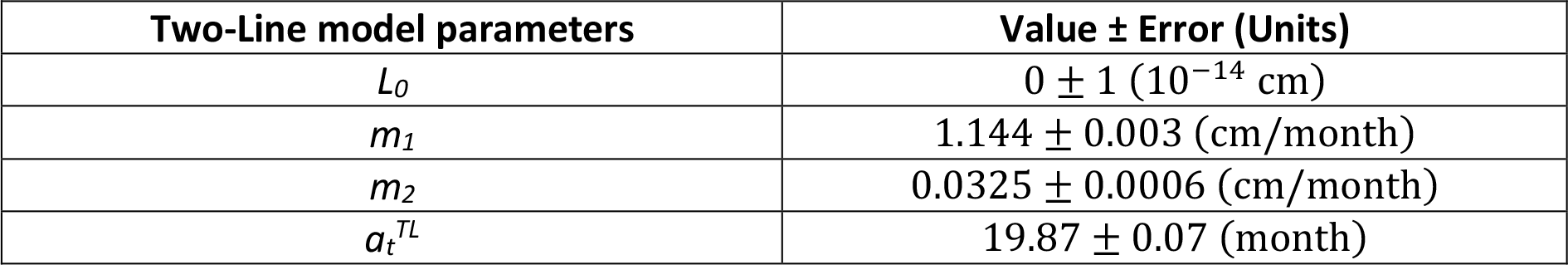
Estimation of the Two-Line model parameters from the normogram of ST lengths.

**Table 3:** Estimation of the Two-Line model parameters from the normogram of SV lengths.

| Two-Line model parameters | Value ± Error (Units) |
| --- | --- |
| $L_0$ | $8 \pm 2$ ($10^{-16}$ cm) |
| $m_1$ | $0.618 \pm 0.002$ (cm/month) |
| $m_2$ | $0.0202 \pm 0.0004$ (cm/month) |
| $a_t^{TL}$ | $19.00 \pm 0.07$ (month) |

### Fitting the growth model to the experimental normograms

In this study, we fitted the following linear growth model to the normograms of ST and SV lengths:

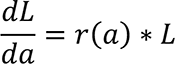

Where, *L* is the ST or SV length at age *a* and *r*(*a*) is the growth rate (depending on the age *a*), defined as follows:

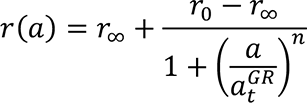

The model free-parameters are the *r_0_*, *r_∞_*, *n* and *a_t_^GR^*, the growth rate at age zero, the growth rate at infinite age, the Hill-exponent denoting sigmoidicity and the growth rate transition age, respectively.

By fitting the model to the experimental data and using bootstrapping, we estimated the parameter values together with their errors from the ST (table 4) and SV (table 5) lengths.

**Table 4:** Estimation of the linear growth model parameters from the normogram of ST lengths.

| Growth model parameters | Value ± Error (Units) |
| --- | --- |
| $r_0$ | $0.323 \pm 0.002$ (1/month) |
| $r_{\infty}$ | $0.00116 \pm 0.00003$ (1/month) |
| $n$ | $5.6 \pm 0.07$ (—) |
| $a_t^{GR}$ | $9.87 \pm 0.05$ (month) |

**Table 5:** Estimation of the linear growth model parameters from the normogram of SV lengths.

| Growth model parameters | Value ± Error (Units) |
| --- | --- |
| $r_0$ | $0.247 \pm 0.001$ (1/month) |
| $r_{\infty}$ | $0.00142 \pm 0.00002$ (1/month) |
| $n$ | $7.8 \pm 0.1$ (—) |
| $a_t^{GR}$ | $11.07 \pm 0.05$ (month) |

Jupyter Notebook (http://jupyter.org/) containing the source code for all computations performed and referred to as Riquelme-Guzmán et al., 2021 in this study can be found at https://doi.org/10.5281/zenodo.5033614. ^51^

### Statistical analysis

Statistical analysis was performed using the software Prism9 (GraphPad Software, LLC) for macOS. To assess the differences in ST or SV between sexes (Fig. 1H), a paired t test was performed between samples. To assess the differences in volume between radii and ulnas of different ST animals (Fig. 2E) and between radii and ulnas of paedomorph and metamorphic animals (Fig. 8C), a Two-way ANOVA test was performed, using a post hoc Tukey’s multiple comparisons test for assessing statistical significance between each sample pair. P-values < 0.05 were considered statistically significant.

## Acknowledgements

We would like to thank Elly M. Tanaka for providing the *Sox9-Tg* line and Joshua D. Currie for guidance during the creation of the *Tie2-Tg* line. We thank Prof. Lorenz Hofbauer, Osvaldo Contreras, Rita Aires and Sean Keeley for proof-reading this paper and for their valuable comments. We thank all members of the Sandoval-Guzmán Lab for continuous support during the development of this work. We also would like to thank Anja Wagner, Beate Gruhl and Judith Konantz for their valuable dedication to the axolotl care. This work was funded by a DFG Research Grant (SA 3349/3-1). CRG was supported by the DIGS-BB Fellow award. AC and OC were funded by Consejo Nacional de Investigaciones Científicas y Técnicas (CONICET) of Argentina and by the grant from Agencia Nacional de Promoción Científica y Tecnológica (ANPCyT) PICT-2017-2307. AC was supported by a doctoral scholarship program from CONICET. OC is a career researcher from CONICET. This work was supported by the Light Microscopy Facility, a Core Facility of the CMCB Technology Platform at TU Dresden.

## Authors contribution

CRG and TSG conceived experiments, analysed data and wrote the manuscript. CRG conducted experiments. MR, MS, AB and SE contributed experimental data and support. AC and OC performed the mathematical modelling. DK created *Sox9-Tg* line. DK, OC and MR contributed with manuscript revision and discussions. TSG secured funding for the project.

